# Differential Gene Expression in the Tropical House Cricket and Its Iridovirus in Healthy versus Diseased Specimens

**DOI:** 10.64898/2026.05.19.726264

**Authors:** Joseph A. Hinton, Hunter K. Walt, Kristin R. Duffield, José L. Ramírez, Florencia Meyer, Federico G. Hoffmann

## Abstract

The tropical house cricket, *Gryllodes sigillatus*, is a mass-produced insect that is used as a protein source for pets and livestock. However, intensive mass-rearing conditions, coupled with high genetic relatedness, create an ideal environment for the spread of pathogenic microbes that severely impact production. Cricket iridovirus (CrIV) is a pathogen that impedes cricket growth and causes significant losses for cricket farmers. Interestingly, recent studies have shown that CrIV is often present asymptomatically, yet the molecular basis of the emergence of disease symptoms remains unknown. To address this, we sampled healthy and diseased crickets and examined differences in cricket and CrIV gene expression via RNAseq. Using differential gene expression analysis and functional enrichment analysis, we found significant differences in host and viral gene expression between healthy and diseased crickets, including genes involved in immunity. Interestingly, while we observed high CrIV gene expression across the entire CrIV genome in sick populations, healthy asymptomatic populations showed elevated expression at a single viral locus. Our results shed light not only on the cricket immune response to CrIV infection but also identify a viral gene that is highly expressed during covert infections, suggesting its potential role in suppressing the host’s immune response. These findings enhance our understanding of how CrIV interacts with its cricket host, providing essential insights for developing targeted strategies to manage CrIV outbreaks in cricket mass-rearing facilities.

## Introduction

As the world pushes to find more sustainable solutions to modern problems, insect farming has emerged as a key player. While mass-reared insects are often used as pet food, they have also recently been put forward as potential solutions to several industrial issues. Black soldier fly (BSF) and yellow mealworms have been proposed as tools to manage waste (Peng et al., 2021; Siddiqui et al., 2022), while the frass from BSF and cricket farming can be used as fertilizer (Lomonaco et al., 2024; Ony et al., 2024). Parasitoid wasps have also been used in agricultural pest management settings (Parra & Coelho, 2022). Current research has even shown that insects can be used to produce biofuel (Koyunoğlu, 2024) and their exoskeletons can be made into bioplastics (Mallegni et al., 2025). Many edible insects are now used as an alternative protein source for livestock and family pets, as well as for human consumption (Cunha et al., 2023; Hancz et al., 2024; T. K. Kim et al., 2019). As the insect industry keeps expanding, continued research is necessary to provide solutions to the major problems plaguing production, including maintaining insect welfare (Barrett & Fischer, 2023), high initial costs, issues at large scale (Biteau et al., 2024), and, in particular, the spread of disease within and between production facilities (Lecocq & Jensen, 2025).

The conditions found in mass rearing facilities, where a large number of animals are kept in close proximity, are conducive to pathogen transmission and often lead to epidemics that result in significant losses. Insect farms are constantly dealing with bacterial, fungal, and viral infections (Akemu et al., 2025; Lecocq & Jensen, 2025). Insect viruses have impacted nearly every corner of the industry, causing grassarie in silkworms (Chopade et al., 2021), chronic bee paralysis in honeybees (Budge et al., 2020), and mass mortality in super worm farms (Penzes et al., 2024). Cricket farms have also been impacted by viral infection, with epidemics of *Acheta domesticus* densovirus (AdDNV), cricket paralysis virus, *Gryllus bimaculatus* nudivirus, and many other cricket viruses devastating the pet feed industry (Huger, 1985; Pham, Bergoin, et al., 2013; Pham, Yu, et al., 2013; Reinganum, 1975; Szelei et al., 2011; Wang et al., 2007). These epidemics can lead to shifts in the cricket-rearing industry towards using alternative cricket species that are resistant to some of these viruses. For instance, the AdDNV-resistant tropical house cricket, *Gryllodes sigillatus*, is another mass-produced cricket that is commonly used as a protein source in feed and food (Magara et al., 2021; Weissman et al., 2012).

While the tropical house crickets are resistant to AdDNV, other viruses still impact them, including cricket iridovirus (CrIV), an isolate of which was described by members of our team (Duffield et al., 2021). CrIV is a large, double-stranded DNA virus of the genus *Iridovirus*. Members of this genus infect many different arthropods and can cause disease (Chinchar et al., 2017). The symptoms of CrIV infection include milky hemolymph, swollen abdomens, and a blueish sheen, which reduce fecundity and increase mortality within impacted cricket colonies. These infections can result in a significant loss of production for the cricket industry. However, CrIV can be present in *G. sigillatus* individuals that do not show any of these symptoms and appear to be healthy (Duffield et al., 2021). It is unknown what determines which individuals show clear signs of infection compared to subclinical infections.

We hypothesize that crickets with signs of disease will have higher expression of genes involved in pathogen defense and immune response. To test this hypothesis, Duffield et al. used qPCR to quantify the expression of fourteen genes related to the cricket immune pathway during overt viral infection (Duffield et al., 2022). They found that the expression of twelve of these genes was significantly higher in crickets with overt infections. In this study, we leverage a recently released chromosome-level assembly of the tropical house cricket genome (Zhang et al., 2025) and use RNA-Seq to characterize genome-wide changes in gene expression associated with disease. Our results revealed multiple genes with consistent changes in immune gene expression across both males and females, dependent on disease state. Interestingly, we also found that CrIV gene expression was different between diseased and healthy crickets and identified a gene that is highly expressed across all samples, including those with subclinical infection.

## Materials and Methods

### Cricket Sampling, RNA Extraction, and RNAseq

Cricket samples used in this study were obtained from lab-reared diseased and healthy *Gryllodes sigillatus* colonies. The source and maintenance of these colonies are described in previous studies (Duffield et al., 2021, 2022). Diseased colonies were characterized by high mortality among late-instar nymphs and adults, strong putrid odor, and milky white hemolymph, which appeared iridescent under illuminated magnification (Duffield et al., 2021). Colonies were maintained under standard cricket rearing protocols, housed in 55L plastic storage bins with ventilated lids packed with egg carton to increase rearing surface area, and housed in an environmental chamber at 32□ on a 16-hour:8-hour light:dark cycle. They were fed a standard diet of Mazuri^®^ Rat and Mouse Diets pellets, Purina^®^ Cat Chow Complete pellets, and water (glass vials plugged with moist cotton) ad libitum. Adults from diseased and healthy colonies were collected seven days after final molting and were killed by freezing at –80□ until used.

Whole-body samples were sent to LC Sciences (Houston, TX) for RNA extraction, library preparation using random hexamers, and RNA sequencing. Briefly, RNA was extracted via TRIzol (Invitrogen, USA) following the manufacturer’s protocol. Total RNA quality and quantity were assessed with Bioanalyzer 2100 and RNA 6000 Nano LabChip kit (Agilent, USA), and all samples obtained RIN numbers greater than 7.0. RNAseq sequencing was conducted using 2×150bp paired-end sequencing (PE150) on an Illumina NovaSeq 6000 following the vendor’s recommended protocol.

### Cricket Gene Expression Analysis

The *Gryllodes sigillatus* genome assembly (GenBank Accession: GCA_965111905.1) was screened for contaminant sequences using ContScout (Bálint et al., 2024) through Docker v25.0.2 (Merkel, 2014). Any sequence that was labelled as outside of the kingdom Metazoa by ContScout was removed from the data by the filterbyname script contained within the BBMap v39.06 suite (Bushnell, 2014). FastQC v0.12.1 (Andrews, 2010) was used to provide reports of the quality of the reads, and the data was then preprocessed using fastp v0.23.4 (S. Chen et al., 2018) to trim low-quality reads and adapter sequences from the data. Next, the processed reads from each sample were quantified with salmon v1.10.3 (Patro et al., 2017) using automatic detection of the library type and the validateMappings flag. The salmon index was created from the combined transcriptome and genome, using the genome as decoys and the gencode flag. The reads were then imported into R v4.4.2 (R Core Team, 2019) using tximport (Soneson et al., 2016). DESeq2 v1.42.1 (Love et al., 2014) was used for differential gene expression analysis. Genes were labelled as significant if they had a Benjamini-Hochberg adjusted *p*-value smaller than 0.05 and a log2FC less than –1 or greater than 1. Gene Ontology (GO) terms were assigned using EggNOG-mapper v2.1.9 (Cantalapiedra et al., 2021) in diamond mode using the EggNOG database v5.0 (Huerta-Cepas et al., 2019). topGO v2.60.1 (Alexa & Rahnenführer, 2007) was then used to identify enriched functions from the differentially expressed genes, using the Fisher statistic and weight01 algorithm. rrvgo v1.16.0 (Sayols, 2023) was used to reduce the GO annotations into a smaller number of categories for easier visualization, with the categories being combined based on semantic similarity. The semantic similarity matrix used to reduce the number of categories was constructed using the *Drosophila melanogaster* organism annotation database, org.Dm.eg.db v3.21 (Carlson, 2019), with the relative distance method and scored as the log_10_ of the *p*-value. It was then reduced using a similarity threshold of 0.95.

### Viral Gene Expression Analysis

Clean reads from each sample were mapped to the CrIV genome (GenBank Accession: GCA_031287495.1, isolate: Liz-CrIV_USDA_2019) using HISAT2 v2.2.1 (D. Kim et al., 2019) with default parameters. The resulting SAM alignment files were converted to binary (BAM) format using samtools v1.6 (Danecek et al., 2021) in “view” mode and sorted and indexed using the “sort” and “index” commands, respectively. Depth across the CrIV genome was calculated using samtools in “depth” mode, using the –aa option to output depth at all positions for each sample across the genome. The average depth/coverage across the CrIV genome was calculated for both the diseased and healthy samples and was then visualized using ggplot2 v3.5.2 (Wickham, 2016) in R.

To determine differences in expression across individual CrIV genes, transcript abundance was estimated using salmon v.0.14.1 and the coding sequence file associated with the CrIV reference genome. Counts were imported to R using tximport and normalized for library size using the trimmed mean of M-values method (TMM) in edgeR v4.4.1 (Y. Chen et al., 2025). Counts per million (CPM) for each gene were calculated using edgeR. The 25 most highly expressed genes were visualized using ComplexHeatmap v2.22.0 (Gu et al., 2016).

To determine if the disease state is caused by coinfection with other viruses, we mapped the reads to the genomes of other common cricket viruses using HISAT2. The reads were mapped to the *Acheta domesticus* densovirus genome (GCF_003033195.1), the *Acheta domesticus* iflavirus genome (GCF_023156815.1), the *Acheta domesticus* mini ambidensovirus genome (KF275669.1), the two segments of the *Acheta domesticus* segmented densovirus genome (OP436269.1, OP436270.1), the *Acheta domesticus* volvovirus genome (KC543331.1), the cricket paralysis virus genome (NC_003924.1), the cricket iridovirus genome (GCA_031287495.1), and the *Gryllus bimaculatus* nudivirus genome (GCF_000870665.1).

## Results

### Transcriptomic Changes due to Disease State by Sex

We checked the publicly available *Gryllodes sigillatus* genome assembly (GCA_965111905.1) for contigs that mapped to any non-Metazoan organism, resulting in 47 contigs being removed from the data. These contigs were found to be microbial contamination, notably with one large scaffold containing a majority of the CrIV genome. We also identified a sample, GS12, which was labelled as a diseased female cricket, but clustered with the healthy female crickets in a transcriptome heatmap and principal component analysis. It also had CrIV expression consistent with a healthy female cricket, which led us to remove the sample from the analysis (Figure S1). In our initial analyses, we found that sex was the main driver of variation in cricket gene expression, accounting for 98% of the transcriptome variation among the samples (Figure 1A). Due to the impact of sex on the variation, we decided to split the analysis by sex. After the separation, we found that the samples began to cluster by disease state, with the first principal component accounting for 70% of the variance in female crickets (Figure 1B) and 46% of the variance in male crickets (Figure 1C).

**Figure 1:**
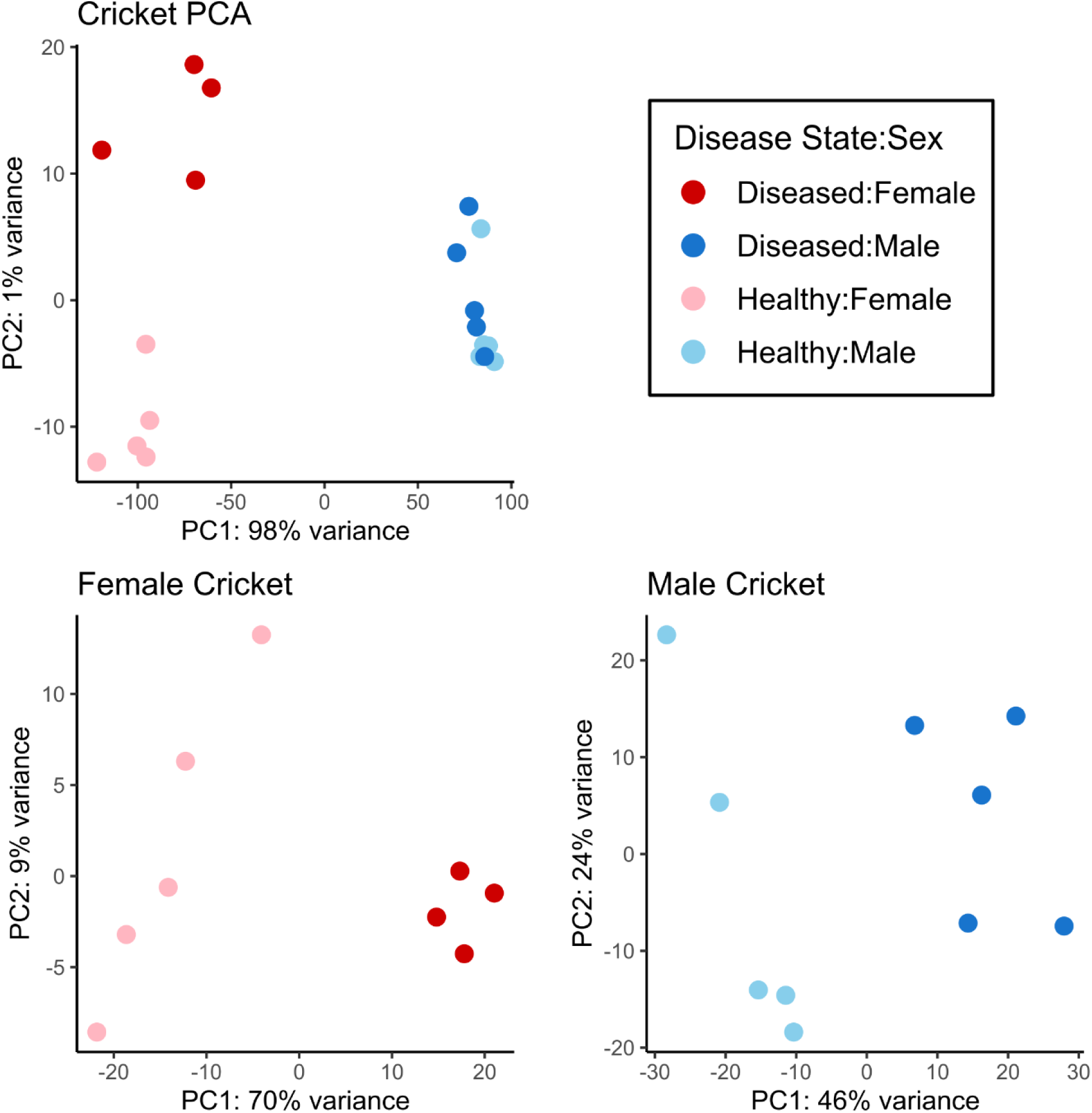
Variation of cricket gene expression due to sex and disease state. (A) PCA plot showing the first two principal components, representing 99% of the variance for combined male and female cricket samples. (B) PCA showing the first two principal component, representing 79% of the variance for female cricket samples. (C) PCA showing the first two principal components, representing 70% of the variance for male cricket samples. Red dots represent female crickets, with lighter dots indicating a healthy sample and darker dots indicating a diseased sample. Blue dots represent male crickets, with lighter dots indicating a healthy sample and darker dots indicating a diseased sample.

When comparing the transcriptomes of males and females separately, we detected the expression of 12,766 genes in male crickets, 741 of which were significantly upregulated and 799 of which were significantly downregulated in diseased crickets. The corresponding numbers were lower for female crickets. We detected the expression of 11,749 annotated genes in female crickets, 598 of which were significantly upregulated and 591 were significantly downregulated in diseased individuals (Figure 2A-B). Gene Ontology (GO) enrichment analysis showed overrepresentation of some immune functions, such as the Toll signaling pathway in male crickets and negative regulation of the innate immune response in female crickets (Figure S2). The GO analysis also showed that functions that were not directly related to immunity, such as cytoplasmic translation and ribosome biogenesis, were overrepresented in both sexes.

**Figure 2:**
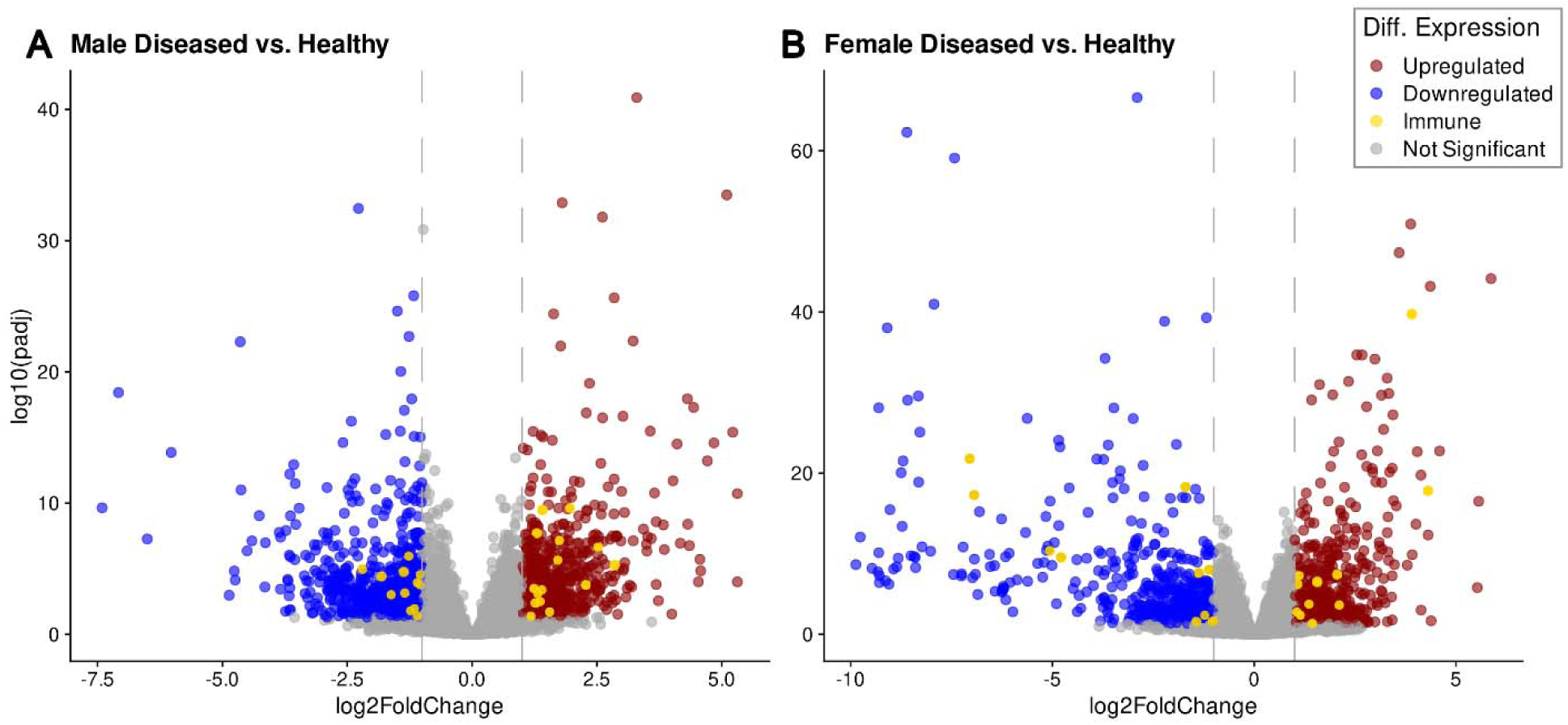
Differentially expressed genes by disease state in male crickets and in female crickets. (A-B) Volcano Plots showing log_2_ of the fold change and the log of the adjusted *p*-value for male (A) and female (B) crickets. Genes with a log_2_FC > 1 and a log(padj) > 1 are considered upregulated. Genes with a log_2_FC < –1 and a log(padj) > 1 are considered downregulated. Differentially expressed genes that were annotated with GO:0002376 (immune system process) from eggNOG are labeled as immune.

### Functional Changes within the Transcriptome due to Disease State

We identified 381 genes that were classified as differentially expressed in both sexes (Figure 3A). Among them, there was overrepresentation of some GO terms associated with immune functions, such as “defense response to Gram-positive bacterium” and “negative regulation of innate immune response” (Figure 3B). To explore whether the expression of immunity-related genes was associated with disease state, we focused on the genes labeled with the GO term “immune system process” (GO:0002376). In the male samples, 27 of the differentially expressed genes were labeled as GO:0002376 (Table S1 and Table S2), whereas 22 of the differentially expressed genes in the female samples were labeled with this GO Term (Table S3 and Table S4). Six immunity-related genes were found to be differentially expressed in both sexes. Five of these genes were upregulated in diseased crickets for both sexes. These genes were a prophenoloxidase gene, a serine protease inhibitor gene, a gene belonging to the PDGF VEGF growth factor family, an integrin gene, and a gene containing a zinc finger domain with no functional description. The sixth gene was upregulated in healthy crickets, or conversely downregulated in diseased crickets, for both sexes and shared high sequence similarity to peptidoglycan recognition proteins in insects.

**Figure 3:**
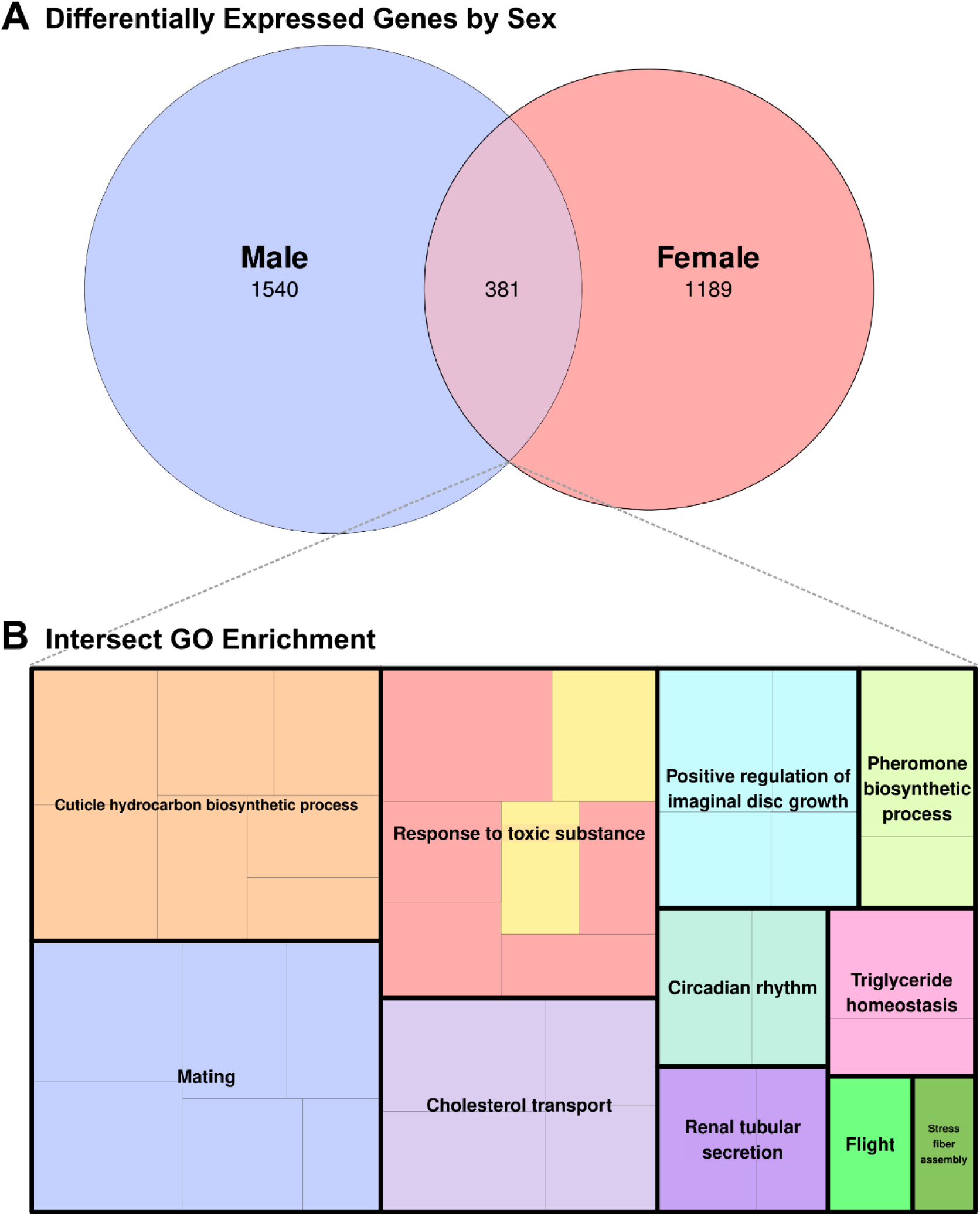
Genes that are differentially expressed in both male and female crickets. (A) Venn Diagram showing the number of differentially expressed genes in male and female crickets and their intersection. (B) Tree map of parent terms derived from functional enrichment analysis of the differentially expressed genes shared by both males and females. The highlighted cells within the “Response to toxic substance” parent term are immune-related (“defense response to Gram-positive bacterium” and “negative regulation of innate immune response”).

### Viral Expression within Diseased and Healthy Crickets

Analysis of the CrIV gene expression within the samples showed that many of the viral genes are highly expressed in the diseased crickets (Figure 4A), where we found coverage across the majority of the 208 predicted genes in the viral genome, with an average coverage of 75.6% when including GS12 and an average of 82.9% when excluding GS12 (Figure 4B). Further statistics and averages will exclude GS12, as the sample was not included for differential expression analysis or any other analyses. In contrast, there appeared to be little to no RNA detected for most viral genes in the healthy crickets, with only 7.5% average coverage across the genome (Figure 4C). Across the healthy crickets, there was a single gene, called 206R, that was highly expressed compared to the rest of the genes. In fact, 206R was highly expressed in nearly every cricket sample, regardless of disease state, but, unfortunately, functional information could not be predicted for this gene. Other genes that were still expressed across most healthy crickets include 393L, 067R, and 030L. The full expression profile is shown in Figure S3, and coverage profiles without the log transformation are shown in Figure S4.

**Figure 4:**
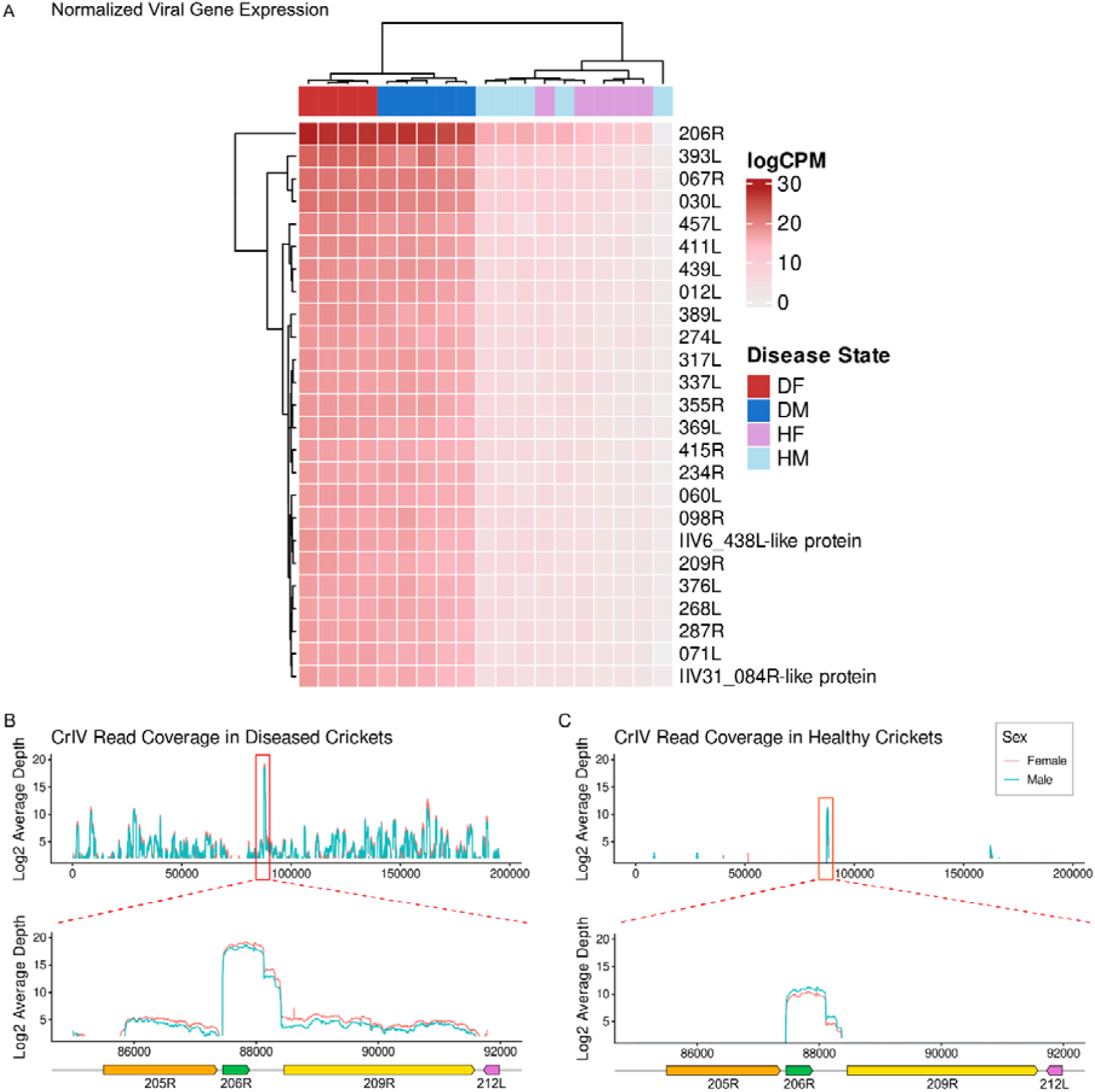
Expression across the CrIV genome between *G. sigillatus* healthy and diseased specimens. (A) Heatmap showing the top 25 most highly expressed CrIV genes (logCPM) across the cricket samples. Samples labeled DF are diseased females, HF are healthy females, DM are diseased males, and HM are healthy males. A separate heatmap showing all viral genes is provided in Supplemental Figure 3 (B) Average read depth from all of the diseased cricket samples across the CrIV genome. The red box indicates the 206R region that has been zoomed in on in the lower panel. (C) Average read depth from all healthy cricket samples across the CrIV genome. The red box indicates the 206R region that has been zoomed in on the lower panel.

Analysis of read mapping to other cricket viruses determined that, if present, these viruses were in very low abundance and there was no clear pattern of differences in abundance between healthy and diseased crickets (Figure S5). The highest number of reads mapped of any non-CrIV virus in any diseased or healthy sample is 392 out of 22,987,151 reads, while the diseased crickets have hundreds of thousands of reads mapping to CrIV and show clear differences in healthy (average 6184 reads) and diseased CrIV abundance (average 1746883 reads). This suggests that viral coinfection does not explain the difference between diseased and healthy crickets.

## Discussion

In our study, we looked for transcriptomic signatures that could explain why some crickets infected with CrIV do not present apparent symptoms, whereas others show clear signs of disease. To do so, we assessed differences in gene expression between healthy and diseased crickets infected with CrIV. We found that sex rather than disease state was the major determinant of differences in gene expression, accounting for most of the variation in gene expression (Figure 1), and as a result, the two sexes were analyzed separately. Our results indicate that differences in disease state are associated with the up– and down-regulation of several hundred genes. The numbers of up– and down-regulated genes were similar for each sex, with disease onset resulting in significant modulation of 7.3 % and 9.5 % of the genes in females and males, respectively. We also found that some functions related to immunity were enriched in the differentially expressed genes, including those involved in the Toll signalling pathway (GO:0008063) and negative regulation of immune response (GO:0045824). Of the six genes labeled with association to the immune system (GO:0002376) that were differentially expressed in both sexes, we found that five were upregulated in diseased crickets, with many of their functions being important for an insect’s immune response. For instance, prophenoloxidase, when activated, produces melanin to encapsulate and destroy pathogens (Cerenius & Söderhäll, 2021). The upregulation of this gene is consistent with a previous study in which black nodules indicating melanization were found in crickets under active infection (Duffield et al., 2021). Serine protease inhibitors (serpins) regulate immune responses by inhibiting serine protease cascades, including those in the prophenoxidase/melanization pathway and in the production of antimicrobial peptides (Meekins et al., 2017). Integrin is a component of hemocytes, which can recognize pathogens and may also regulate immune proteins in fat body cells (Kausar et al., 2022). The single immune gene that was upregulated in healthy crickets across both sexes encodes a PGRP-LB-like protein. Previous studies have shown that PGRP-LB and similar genes have an antiviral function against other insect viruses (Y. Chen et al., 2015; Li et al., 2023). The increased expression of this gene could prevent overt CrIV infection in the healthy population. The immune genes that are upregulated in diseased crickets, with the exception of the prophenoloxidase gene, do not directly combat viral particles, but instead signal the presence of viral particles. Our results are not consistent with a previous study, which found clear upregulation of immune signaling genes using qPCR (Duffield et al., 2022). The discrepancies may be due to the increased sensitivity of qPCR compared to RNA sequencing in this study. Additionally, whole-body transcriptome analysis rather than a subset of immune genes or specific immune tissues could make specific trends in immune response difficult to identify. Investigating specific tissues like the fat body could be the focus of future studies.

When examining the expression of viral genes, we found clear differences between healthy and diseased crickets, with no apparent differences between sexes. Since there was no clear pattern of expression of other viruses corresponding to disease state, we ruled out coinfection as a contributor to disease (Figure S5). The expression of the other viruses appears to be a mixture of mis-mapping reads (some reads map to the poly-A tail of AdIV) and low-coverage read assignment, limiting detection power. However, the number of reads mapping to CrIV is orders of magnitude higher than for all other viruses. CrIV gene expression is higher in diseased crickets, with coverage across the majority of the genome (Table S5). In contrast, CrIV in healthy crickets appears to express only a small portion of the genome. There were a few viral genes that were still clearly expressed in healthy crickets, but the 206R gene stands out as it is expressed at such a high level. In a previous study, it was shown that the 206R protein contains a predicted functional domain that is conserved in multiple other arthropod viruses, including Lake Sinai virus, cypoviruses, and tospoviruses (Karlin, 2024). The study proposes that the domain, which was named a WIV (widespread, intriguing, and versatile) domain, acts as a virulence factor in arthropod infection. Based on these factors and 206R’s high expression in both active and covert infections of CrIV, it is possible that the 206R protein works to keep the virus hidden from the host immune response. Latency or dormancy is a common strategy that many viruses use to persist long-term within their hosts and evade the host immune system. The sharp expression of a single viral gene in healthy individuals is striking and could be related to the maintenance of the dormant state. It is possible that CrIV infects most crickets while the 206R gene product, alone or in combination with other expressed viral or host genes, is responsible for restricting the expression of other CrIV genes, apoptosis, or one or more host immune pathways, which would allow the virus to persist in a subclinical state. 206R was the highest expressed CrIV gene on average, which made it of particular interest to us. However, 206R is not the only gene that is expressed in healthy crickets. Other genes may play a role in facilitating CrIV covert infection, but sequencing depth was lacking across the CrIV genome in healthy samples (Table S5). The diseased phenotype could also be related to virus load, with disease manifesting beyond a given threshold. Based on the coverage plots, it appears that only a few genes are expressed. However, other genes may have been expressed but were not detected due to the low sequencing depth used for this study, which was not optimal for detecting full viral genomes. This could mean that CrIV exhibits a similar expression profile in healthy crickets, but it is scaled down in comparison to crickets with active infection. Either way, 206R is still highly expressed in both healthy and diseased crickets.

When we investigated the potential function of 206R, we found that the 206R gene is present in other iridoviruses, but that any functional information is not currently available (Bronkhorst et al., 2012; de Faria et al., 2022). However, de Faria et al (2022) shows that, in *Drosophila melanogaster*, the 206R gene of Invertebrate iridescent virus 6 (IIV-6) is transcribed via the host’s RNA Polymerase II. The resulting dsRNA fragments then trigger the host’s RNAi pathway, releasing siRNAs, small RNAs that target viral genes in the RNAi antiviral response. The 206R gene is shown to be a major target of these siRNAs in the study. CrIV may be recognized by a similar mechanism in crickets, meaning the expression of the RNAi pathway or of siRNAs could be a determining factor in overt infections. We identified an Argonaute gene (GSIG20043), part of the RNAi pathway, that was differentially expressed in both male and female crickets, but was upregulated in females and downregulated in males, suggesting a possible sex specific response. Unfortunately, because our study used mRNA sequencing, we were not able to detect siRNAs. Future studies should investigate the small RNA repertoire of healthy versus diseased crickets to determine if the RNAi antiviral response is playing a role in suppressing CrIV.

Although active infection strongly impacts the cricket transcriptome, we only detected a few immunity-related modulated genes. We found a prophenoloxidase gene, a serpin-like gene, an integrin-like gene, and two other immune-related genes that were upregulated in diseased crickets and only a single immune gene, a PRGP-LB-like gene, that was upregulated in healthy crickets. Exploring these five genes and their connection to viral infection may reveal what triggers symptoms in CrIV-infected crickets. The difference may also be due to changes in expression in proteins that are not labeled with GO:0002376, since these genes were not focused on in this study. Since many potential immune genes may be present but were not labeled with the GO term during functional annotation, some manually curated immune gene candidates are highlighted in the tables of differentially expressed genes provided in the Supplementary Information. Alternatively, expression of viral genes such as 206R and 393L or the interaction between the host and these viral proteins may cause the observed phenotypes. Future work should focus on the RNAi antiviral pathway in crickets to determine if the host’s siRNA expression plays a role in preventing active infection. Finally, exploring viral factors contributing to active infection could be a promising future direction, such as functional studies of the 206R gene, using more sensitive methods, like qPCR, to determine the impact of specific genes, or nucleotide polymorphism analyses across the CrIV and *G. sigillatus* genome in covertly versus actively infected crickets.

## Data Availability Statement

RNAseq data generated for this study will be released to NCBI’s Sequence Read Archive under Bioproject PRJNA1541515 upon acceptance. All files associated with genome annotation are available online on Zenodo at https://doi.org/10.5281/zenodo.23192433 as Supplemental Information. Total read count and other information for each sample can be found in Table S6.

## Author Contributions

Conceptualization – JAH, HKW, KRD, FM, FGH, JLR; Data Curation – JAH, HKW, KRD, FGH, JLR; Formal Analysis – JAH, HKW, KRD, FGH, JLR; Funding Acquisition – KRD, JLR; Investigation – JAH, HKW, KRD, FGH, JLR; Methodology – JAH, HKW, KRD, FGH, JLR; Project Administration – HKW, KRD, FGH, JLR; Resources – KRD, FGH, JLR; Supervision – HWK, FGH; Validation – JAH, HKW, FGH; Writing – Original Draft – JH; Writing – Review & Editing – JAH, HKW, KRD, FM, FGH, JLR.

