## Supplemental Tables and Figures for "Differential Gene Expression in the Tropical House Cricket and Its Iridovirus in Healthy versus Diseased Specimens"

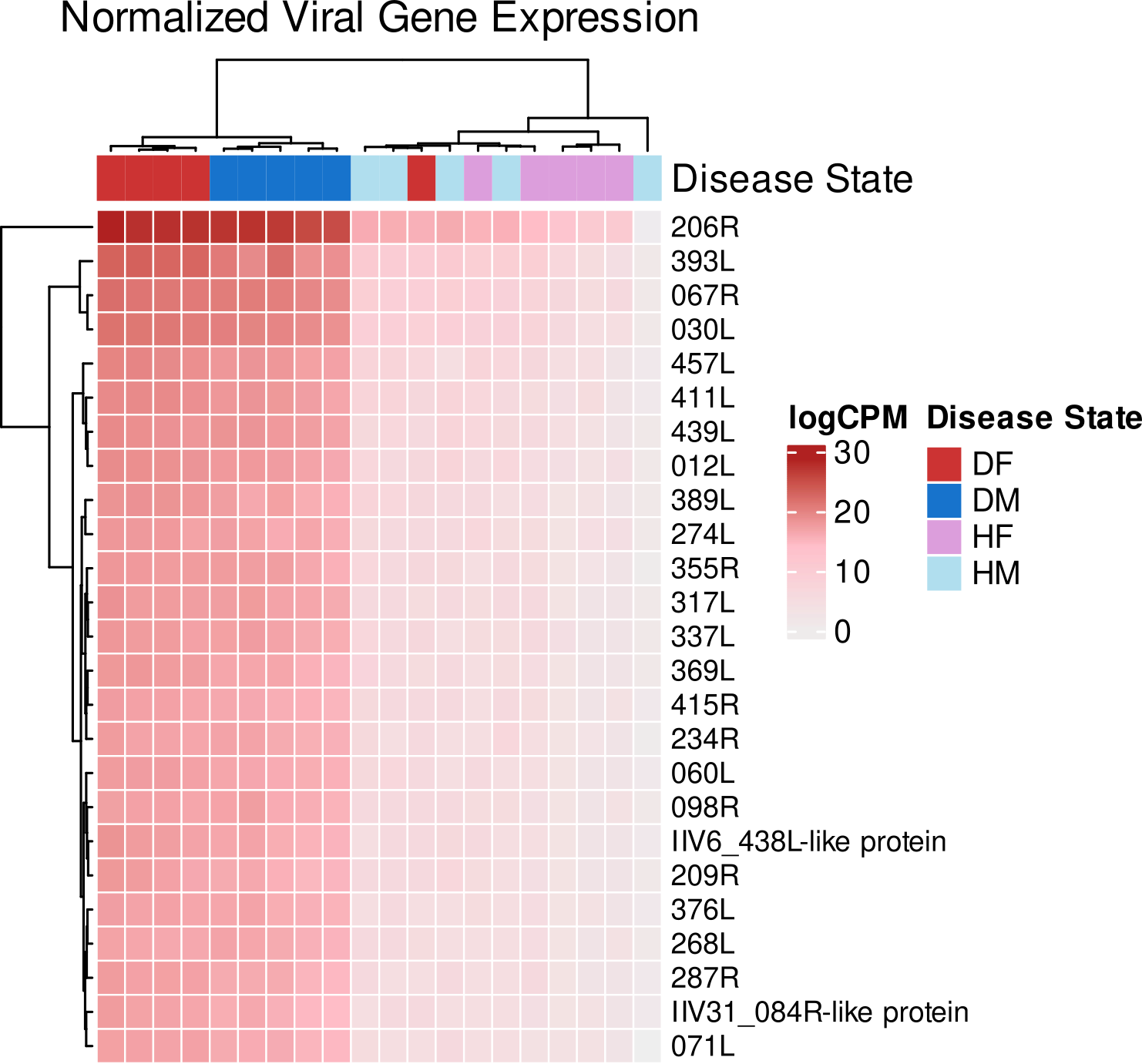

**Figure S1: Viral Expression of All Samples, including a Healthy Cricket from the Diseased Colony**

A heatmap of viral gene expression across all samples in the analysis. GS12, the red sample, is labelled as a diseased female, but clusters with the healthy crickets when comparing viral gene expression.

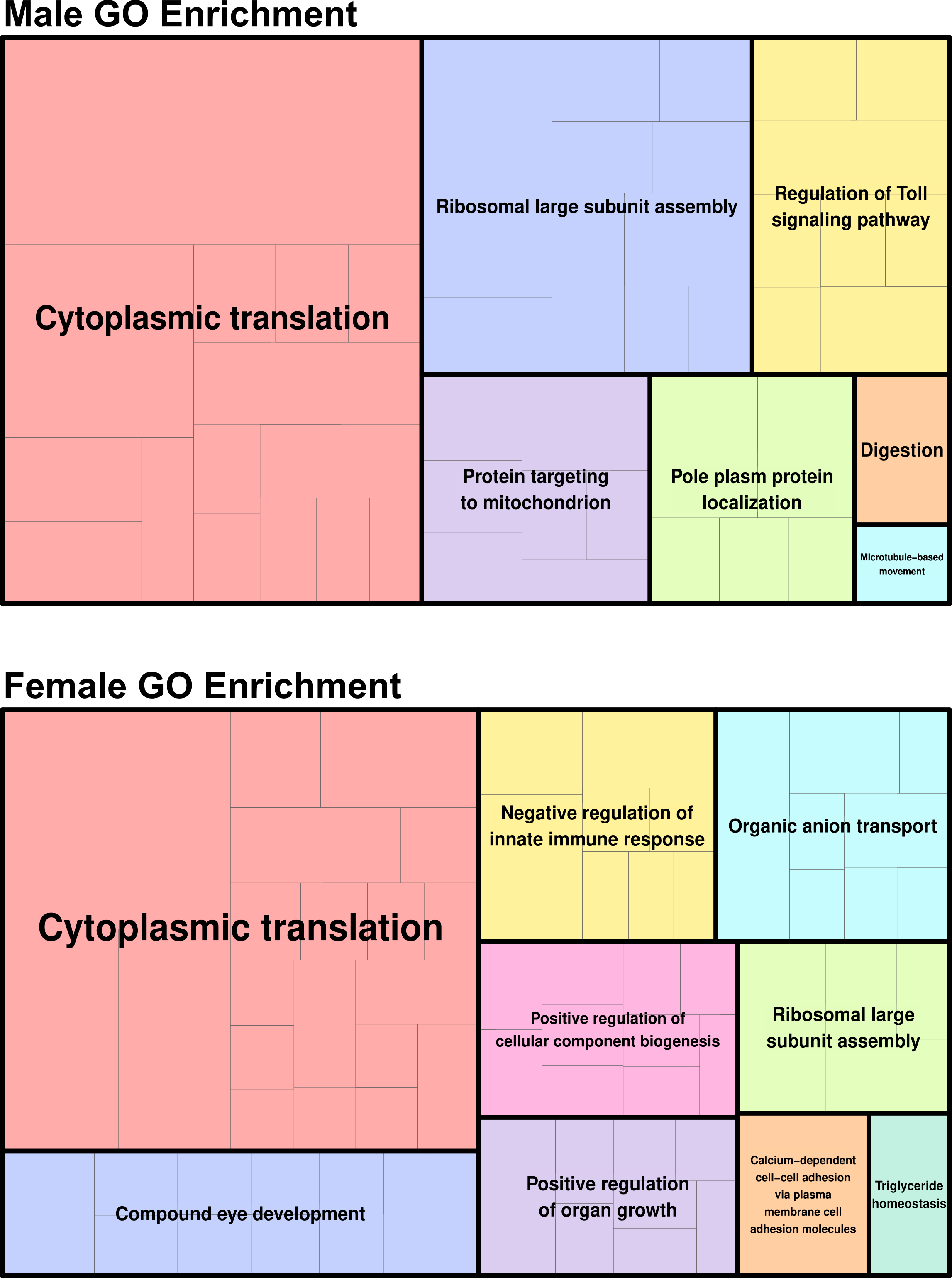

**Figure S2: Functions of Differentially Expressed Genes in Male and Female Crickets**

Treemap plots showing parent terms for the enriched biological process GO terms found in either the differentially expressed genes between male (top) or female (bottom) diseased vs. healthy crickets.

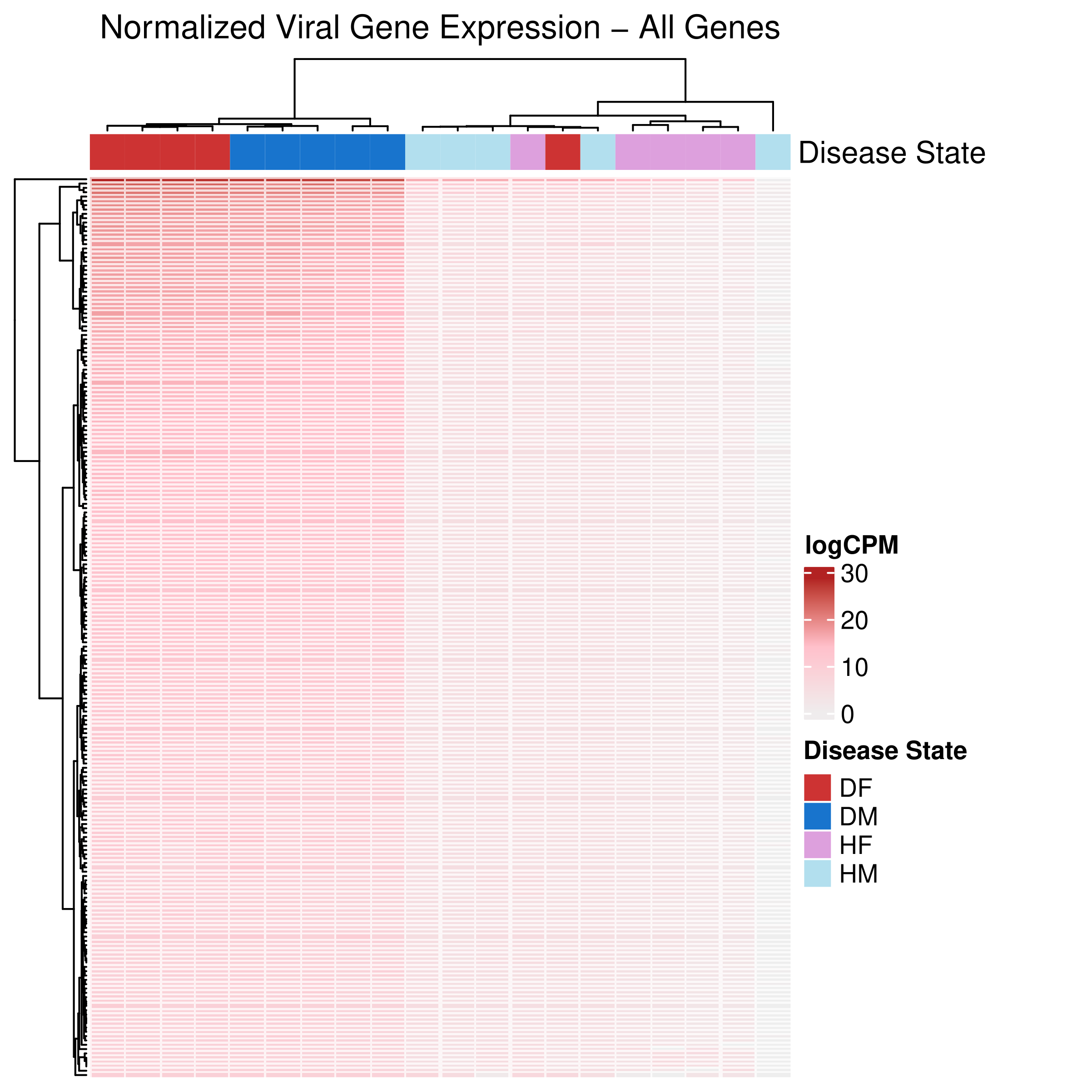

**Figure S3: Expression across the CrIV genome between *G. sigillatus* healthy and diseased specimens (All Genes)**

(A) Heatmap showing all CrIV genes (logCPM) across the cricket samples. Samples labeled DF are diseased females, HF are healthy females, DM are diseased males, and HM are healthy males.

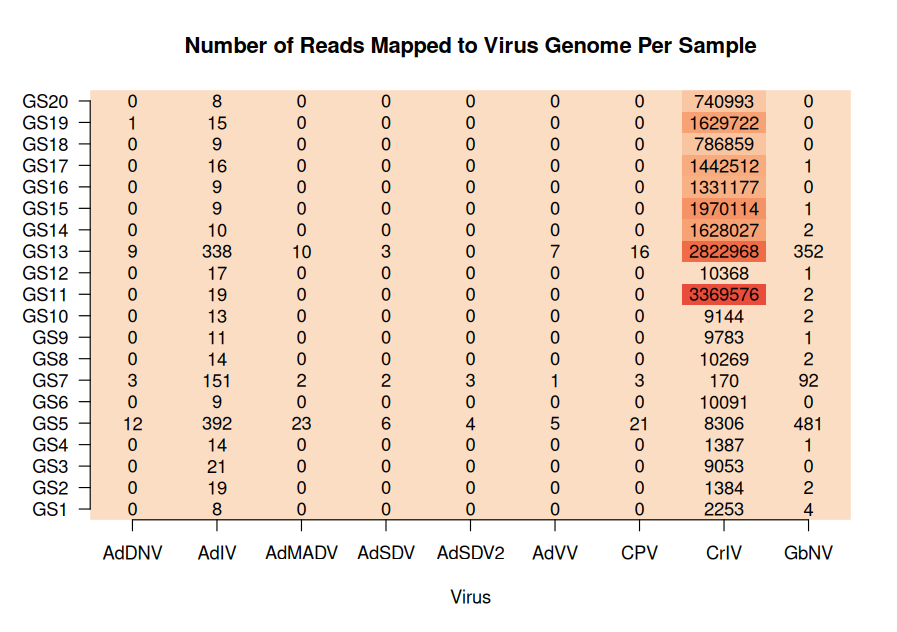

**Figure S4: Reads Mapping to Common Cricket Viruses in Each Sample**

A heatmap showing how many reads within each sample mapped to common viruses that infect crickets. Very few reads map to any viruses, and no clear patterns of expression are seen between healthy and diseased samples.

**Table S1**

Differentially Expressed Genes that are Up-regulated and are associated with Immunity (GO:0002376) in Male Crickets

| **Genes** | **Base Mean** | **log2FoldChange** | **padj** | **Description*** |
| --- | --- | --- | --- | --- |
| GSIG04501 | 165.3302261 | 1.315501962 | 0.000791 | Hydrolase activity, hydrolyzing O-glycosyl compounds. It is involved in the biological process described with carbohydrate metabolic process |
| GSIG06375 | 31.34071869 | 1.956717267 | 2.44E-10 | serine-type endopeptidase activity. It is involved in the biological process described with proteolysis |
| GSIG06414 | 43.23672486 | 1.751663756 | 6.88E-08 | It is involved in the biological process described with signal transduction |
| GSIG06868 | 34.71816968 | 1.716670043 | 2.25E-06 | serine-type endopeptidase activity. It is involved in the biological process described with proteolysis |
| GSIG06869 | 57.96111755 | 1.362900423 | 0.003122 | Domain abundant in complement control proteins; SUSHI repeat; short complement-like repeat (SCR) |
| GSIG06873 | 24.01308927 | 1.553025435 | 0.019313 | serine-type endopeptidase activity. It is involved in the biological process described with proteolysis |
| GSIG07589 | 25.21988081 | 2.518040134 | 2.26E-07 | Belongs to the PDGF VEGF growth factor family |
| GSIG14237 | 51.20271348 | 1.179144184 | 0.0407 | Belongs to the serpin family |
| GSIG18555 | 36.38876062 | 2.859440915 | 5.24E-06 | Oxidoreductase activity. It is involved in the biological process described with metabolic process |
| GSIG18890 | 171.4815858 | 1.299325307 | 1.89E-08 | TRAF6 |
| GSIG19632 | 158.1680822 | 1.409265195 | 3.38E-10 | Common central domain of tyrosinase |
| GSIG21079 | 22.56780029 | 2.280800079 | 0.000163 | Belongs to the serpin family |
| GSIG21136 | 18.76104176 | 1.396542358 | 0.000408 | Integrin beta |
| GSIG22498 | 199.899492 | 1.251643591 | 0.000338 | Broad-Complex, Tramtrack and Bric a brac |
| GSIG24488 | 26.51798254 | 1.26242175 | 0.003985 | Broad-Complex, Tramtrack and Bric a brac |

*Descriptions are assigned by eggNOG-mapper

**Table S2**

Differentially Expressed Genes that are Down-regulated and are associated with Immunity (GO:0002376) in Male Crickets

| **Genes** | **Base Mean** | **log2FoldChange** | **padj** | **Description*** |
| --- | --- | --- | --- | --- |
| GSIG00322 | 22.96687 | -1.23145 | 0.015551 | CLASP N terminal |
| GSIG02367 | 157.5161 | -1.03799 | 2.82E-05 | Phosphatidylinositol 3-kinase regulatory subunit P85 inter-SH2 domain |
| GSIG03803 | 41.99144 | -1.04524 | 0.000163 | Belongs to the DOCK family |
| GSIG07693 | 454.2802 | -1.26643 | 1.08E-06 | It is involved in the biological process described with signal transduction |
| GSIG08182 | 574.5663 | -2.19563 | 1.11E-05 | N-acetylmuramoyl-L-alanine amidase |
| GSIG14240 | 253.9488 | -1.09744 | 0.000115 | Belongs to the serpin family |
| GSIG14250 | 33.8161 | -1.62195 | 0.000944 | Belongs to the serpin family |
| GSIG21399 | 204.0104 | -1.15446 | 0.010932 | It is involved in the biological process described with protein phosphorylation |
| GSIG23698 | 25.46008 | -1.34788 | 0.000715 | Broad-Complex, Tramtrack and Bric a brac |
| GSIG24446 | 15.80438 | -1.81264 | 4.00E-05 | Broad-Complex, Tramtrack and Bric a brac |
| GSIG24451 | 72.90417 | -1.09564 | 0.038638 | Broad-Complex, Tramtrack and Bric a brac |
| GSIG26113 | 170.0735 | -1.37116 | 1.82E-05 | Protein of unknown function (DUF1154) |

*Descriptions are assigned by eggNOG-mapper

**Table S3**

Differentially Expressed Genes that are Up-regulated and are associated with Immunity (GO:0002376) in Female Crickets

| **Genes** | **Base Mean** | **log2FoldChange** | **padj** | **Description*** |
| --- | --- | --- | --- | --- |
| GSIG06414 | 174.3635 | 1.055525 | 0.001741 | It is involved in the biological process described with signal transduction |
| GSIG07589 | 72.55309 | 3.902502 | 1.90E-40 | Belongs to the PDGF VEGF growth factor family |
| GSIG11600 | 54.85497 | 1.3586 | 0.000177 | Aldose 1-epimerase |
| GSIG12055 | 370.7476 | 1.143207 | 0.004258 | chitin binding. It is involved in the biological process described with chitin metabolic process |
| GSIG18022 | 19059.66 | 1.11093 | 5.37E-08 | Stores iron in a soluble, non-toxic, readily available form. Important for iron homeostasis. Iron is taken up in the ferrous form and deposited as ferric hydroxides after oxidation |
| GSIG18044 | 134.8746 | 1.082178 | 5.85E-07 | receptor activity. It is involved in the biological process described with multicellular organismal development |
| GSIG18555 | 103.3099 | 2.045814 | 4.03E-08 | Oxidoreductase activity. It is involved in the biological process described with metabolic process |
| GSIG21079 | 185.9607 | 4.304667 | 1.49E-18 | Belongs to the serpin family |
| GSIG21136 | 44.05226 | 2.105338 | 0.000229 | Integrin beta |
| GSIG24488 | 26.54989 | 1.439507 | 0.044411 | Broad-Complex, Tramtrack and Bric a brac |
| GSIG26973 | 1614.077 | 1.561064 | 3.08E-07 | ML domain |

*Descriptions are assigned by eggNOG-mapper

**Table S4**

Differentially Expressed Genes that are Down-regulated and are associated with Immunity (GO:0002376) in Female Crickets

| **Genes** | **baseMean** | **log2FoldChange** | **padj** | **Description*** |
| --- | --- | --- | --- | --- |
| GSIG01151 | 455.0228 | -1.11668 | 9.42E-09 | Receptor binding. It is involved in the biological process described with regulation of cell migration |
| GSIG02505 | 145.2509 | -1.00569 | 0.021429 | Belongs to the ubiquitin-conjugating enzyme family |
| GSIG08182 | 1251.199 | -4.78691 | 2.81E-10 | N-acetylmuramoyl-L-alanine amidase |
| GSIG08183 | 4785.519 | -6.93818 | 5.14E-18 | N-acetylmuramoyl-L-alanine amidase |
| GSIG08889 | 693.2798 | -1.70563 | 5.05E-19 | Calcium ion binding. It is involved in the biological process described with homophilic cell adhesion via plasma membrane adhesion molecules |
| GSIG15333 | 383.9724 | -1.22441 | 0.004304 | Belongs to the serpin family |
| GSIG20442 | 304.5746 | -5.07141 | 4.04E-11 | Common central domain of tyrosinase |
| GSIG20443 | 1845.014 | -7.05166 | 1.58E-22 | Common central domain of tyrosinase |
| GSIG21109 | 209.8663 | -1.3731 | 2.37E-08 | Fibroblast growth factor receptor |
| GSIG22338 | 19.91307 | -1.04343 | 0.025576 | tyrosine serine threonine phosphatase activity. It is involved in the biological process described with protein dephosphorylation |
| GSIG26306 | 52.06828 | -1.428 | 0.028304 | Metal ion binding |

*Descriptions are assigned by eggNOG-mapper

**Table S5**

Viral Coverage Statistics for Each Sample

| **Sample Name** | **# of Reads** | **Bases Covered** | **Coverage** | **Mean Depth** |
| --- | --- | --- | --- | --- |
| GS1 | 2253 | 9296 | 4.8 | 1.7 |
| GS2 | 1384 | 7568 | 3.9 | 1.0 |
| GS3 | 9053 | 18635 | 9.5 | 6.8 |
| GS4 | 1387 | 8611 | 4.4 | 1.0 |
| GS5 | 8306 | 25531 | 13.1 | 5.6 |
| GS6 | 10091 | 20538 | 10.5 | 7.6 |
| GS7 | 170 | 2232 | 1.1 | 0.0 |
| GS8 | 10269 | 19224 | 9.8 | 7.8 |
| GS9 | 9783 | 20065 | 10.3 | 7.4 |
| GS10 | 9144 | 15609 | 8.0 | 6.9 |
| GS11 | 3369576 | 169398 | 86.7 | 2469.4 |
| GS12 | 10368 | 19721 | 10.1 | 7.8 |
| GS13 | 2822968 | 174148 | 89.1 | 2099.4 |
| GS14 | 1628027 | 162875 | 83.3 | 1234.7 |
| GS15 | 1970114 | 166683 | 85.3 | 1493.1 |
| GS16 | 1331177 | 161280 | 82.5 | 1008.8 |
| GS17 | 1442512 | 166276 | 85.1 | 1093.9 |
| GS18 | 786859 | 146823 | 75.1 | 596.1 |
| GS19 | 1629722 | 156234 | 79.9 | 1235.6 |
| GS20 | 740993 | 153598 | 78.6 | 561.7 |

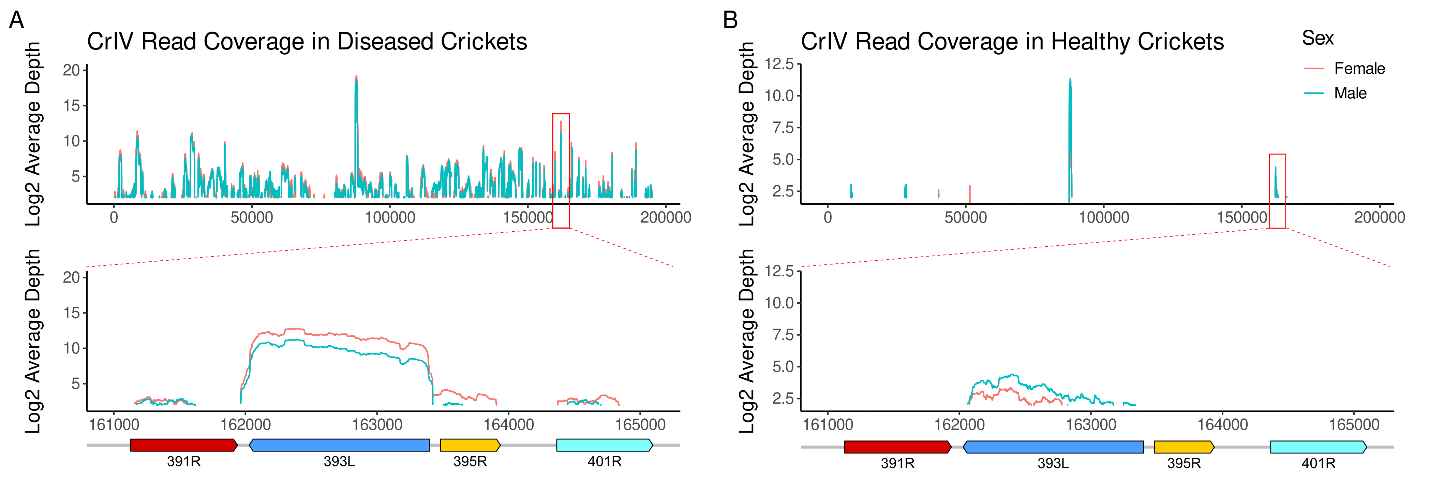

**Figure S6: Viral Expression of the 393L Gene in Healthy and Diseased Crickets**

(A) Average read depth from all of the diseased cricket samples across the CrIV genome. The red box indicates the 393L region that has been zoomed in on in the lower panel. (B) Average read depth from all healthy cricket samples across the CrIV genome. The red box indicates the 393L region that has been zoomed in on in the lower panel.

**Table S6**

| **Sample** | **Total Reads** | **Total CrIV Reads** | **% CrIV Reads** | **Disease State** | **Colony** | **Sex** |
| --- | --- | --- | --- | --- | --- | --- |
| KD1 | 18386975 | 2253 | 0.012253239 | Healthy | Healthy | Female |
| KD2 | 20978399 | 1384 | 0.006597262 | Healthy | Healthy | Female |
| KD3 | 22174824 | 9053 | 0.040825578 | Healthy | Healthy | Female |
| KD4 | 19508224 | 1387 | 0.007109822 | Healthy | Healthy | Female |
| KD5 | 23443789 | 8306 | 0.035429427 | Healthy | Healthy | Female |
| KD6 | 18516338 | 10091 | 0.054497817 | Healthy | Healthy | Male |
| KD7 | 18263702 | 170 | 0.000930808 | Healthy | Healthy | Male |
| KD8 | 21228920 | 10269 | 0.048372692 | Healthy | Healthy | Male |
| KD9 | 19888730 | 9783 | 0.049188661 | Healthy | Healthy | Male |
| KD10 | 22444119 | 9144 | 0.040741185 | Healthy | Healthy | Male |
| KD11 | 24738113 | 3369576 | 13.62099041 | Diseased | Diseased | Female |
| KD12 | 22610618 | 10368 | 0.045854563 | Not Used* | Diseased | Female |
| KD13 | 23583589 | 2822968 | 11.97005256 | Diseased | Diseased | Female |
| KD14 | 17800041 | 1628027 | 9.146198034 | Diseased | Diseased | Female |
| KD15 | 19358378 | 1970114 | 10.17706132 | Diseased | Diseased | Female |
| KD16 | 20265039 | 1331177 | 6.568835125 | Diseased | Diseased | Male |
| KD17 | 18651884 | 1442512 | 7.733867528 | Diseased | Diseased | Male |
| KD18 | 21523442 | 786859 | 3.655823265 | Diseased | Diseased | Male |
| KD19 | 24382083 | 1629722 | 6.684096679 | Diseased | Diseased | Male |
| KD20 | 24069782 | 740993 | 3.078519781 | Diseased | Diseased | Male |

Read Statistics and Metadata for Each Sample
